# Conservative and disruptive modes of adolescent change in brain functional connectivity

**DOI:** 10.1101/604843

**Authors:** František Váša, Rafael Romero-Garcia, Manfred G. Kitzbichler, Jakob Seidlitz, Kirstie J. Whitaker, Matilde M. Vaghi, Prantik Kundu, Ameera X. Patel, Peter Fonagy, Raymond J. Dolan, Peter B. Jones, Ian M. Goodyer, the NSPN Consortium, Petra E. Vértes, Edward T. Bullmore

**Affiliations:** Department of Psychiatry, University of Cambridge, Cambridge, CB2 0SZ, UK; Developmental Neurogenomics Unit, National Institute of Mental Health, Bethesda, MD 20892, USA; The Alan Turing Institute, London, NW1 2DB, UK; Wellcome Trust Centre for Neuroimaging, UCL Institute of Neurology, University College London, London WC1N 3BG, UK; Brain Imaging Center, Icahn School of Medicine at Mount Sinai, New York, NY 10029, USA; Research Department of Clinical, Educational and Health Psychology, University College London, London WC1E 6BT, UK; Max Planck UCL Centre for Computational Psychiatry and Ageing Research, University College London, London WC1B 5EH, UK; School of Mathematical Sciences, Queen Mary University of London, London E1 4NS, UK; Cambridgeshire and Peterborough NHS Foundation Trust, Huntingdon, PE29 3RJ, UK; GlaxoSmithKline R&D, Stevenage SG1 2NY, UK

**Author notes:** P.E.V. and E.T.B. are joint last authors. Email address (František Váša).

**Keywords:** neurodevelopment, connectome, MRI, Allen Human Brain Atlas

## Abstract

Adolescent changes in human brain function are not entirely understood. Here we used multi-echo functional magnetic resonance imaging (fMRI) to measure developmental change in functional connectivity (FC) of resting-state oscillations between pairs of 330 cortical regions and 16 subcortical regions in N=298 healthy adolescents. Participants were aged 14-26 years and were scanned on two or more occasions at least 6 months apart. We found two distinct modes of age-related change in FC: “conservative” and “disruptive”. Conservative development was characteristic of primary cortex, which was strongly connected at 14 years and became even more connected in the period 14-26 years. Disruptive development was characteristic of association cortex, hippocampus and amygdala, which were not strongly connected at 14 years but became more strongly connected during adolescence. We defined the maturational index (MI) as the signed coefficient of the linear relationship between baseline FC (at 14 years, *FC*_14_) and adolescent change in FC (∆*FC*_14−26_). Disruptive systems (with negative MI) were functionally specialised for social cognition and autobiographical memory and were significantly co-located with prior maps of aerobic glycolysis (AG), AG-related gene expression, post-natal expansion of cortical surface area, and adolescent shrinkage of cortical depth. We conclude that human brain organization is disrupted during adolescence by the emergence of strong functional connectivity of subcortical nuclei and association cortical areas, representing metabolically expensive re-modelling of synaptic connectivity between brain regions that were not strongly connected in childhood. We suggest that this re-modelling process may support emergence of social skills and self-awareness during healthy human adolescence.

## 1. Introduction

During adolescence the human brain undergoes substantial changes in both structure (Whitaker et al., 2016; Váša et al., 2017) and function (Paus et al., 2008; Vertes and Bullmore, 2015). Accurately describing these maturational processes is key to understanding the parallel changes in cognition and behaviour, as well as the vulnerability to mental health disorders, that characterize this critical developmental period.

Over the last decade, functional brain networks derived from fMRI have been a useful tool to understand large-scale brain organization (Bullmore and Sporns, 2009; Fornito et al., 2016). The nodes of these fMRI networks correspond to macroscopic brain regions and the edges correspond to the correlations in brain activity, or so-called functional connectivity (FC), between pairs of regionally localised, low frequency oscillations. The architecture of functional networks has been shown to be heritable, affected by both disease and medication, and predictive of treatment outcome (Bullmore and Sporns, 2009). A number of studies have endeavoured to describe adolescent development of functional brain network organisation, but the findings so far have been somewhat inconsistent. This is likely due in part to small sample sizes, the lack of longitudinal data, and significant variation in fMRI data pre-processing and analysis methods (see Table S1 in SI). In addition, although subcortical nuclei are theoretically well-recognised components of frontal cortico-striato-thalamic circuits, subcortical connectivity has generally been measured only for a few nuclei or ignored altogether (see Table S2 in SI).

The most replicable result of early studies of resting-state FC development was an observed increase in the strength of long-range connections accompanied by a decrease in the strength of short-range connections (Fair et al., 2007, 2009; Supekar et al., 2009; Dosenbach et al., 2010). Since long-range connections tend to emanate from association cortical areas involved in higher-order cognitive functions, these results were consistent with prior work suggesting that primary sensory and motor areas mature earlier in childhood, whereas association areas show relatively protracted maturation, extending into adolescence and early adulthood (Mills et al., 2014; Marek et al., 2015; Whitaker et al., 2016; Goyal et al., 2014).

However, it has become clear over the last five years that even the most replicated early findings on developmental changes in FC might have been confounded by the effects of in-scanner head motion (Power et al., 2012; Satterthwaite et al., 2012, 2013; Marek et al., 2015). It is now well-recognised that small (<1 mm), transient head movements during scannning can bias estimation of correlations between fMRI time series and this is a critical issue for developmental studies because younger participants may find it more difficult to remain stationary for several minutes in the scanner.

Here, in an attempt to address the heterogeneity of findings in the literature, we measured resting-state FC maturation in an accelerated longitudinal study of 298 healthy adolescents, aged 14-26 years, each scanned at least twice. To adequately correct fMRI time series for effects of participant in-scanner motion, we used multi-echo scans (Barth et al., 1999) denoised using multi-echo ICA (ME-ICA; Kundu et al., 2012, 2013), a method designed to identify and discard noisy components of fMRI time series unrelated to the BOLD signal. For each of 520 pre-processed and quality-controlled fMRI datasets, we estimated the regional mean time series for each of 330 cortical areas and 16 subcortical nuclei, and the Pearson’s correlation for each possible pair of regions. We used linear mixed effects modelling of repeated fMRI scans on the same participants to explore the hypothesis that there are age-related changes in functional connectivity (FC) of the human brain during adolescence. We identified two modes of developmental change in fMRI connectivity, defined by positive or negative maturational index (MI), and assessed the psychological and biological relevance of these so-called conservative or disruptive systems by meta-analysis of prior task-related fMRI data and by testing for anatomical co-location of the MI map derived from human developmental fMRI with prior maps of cortical anatomy, metabolism, and gene expression.

## Results

### Age-related change of connectivity strengths

The functional connectivity (FC), or correlation between fMRI time series, for a pair of brain regions was generally positive, and there was a weak but significant trend for the global mean correlation to increase with age (see Fig S3, *t*(221) = 2.1, *P* = 0.035). For each regional node, we estimated its strength of connectivity, or equivalently its weighted degree, by averaging the correlations between it and all other brain regions. We also calculated the strength of connectivity specifically within or between cortical and subcortical subsets of nodes. Using a mixed effect linear model of age-related change in these metrics, we estimated the “baseline” strength of FC at age 14 years, *FC*_14_, and the linear rate of change in weighted degree as a function of increasing age, ∆*FC*_14−26_ (Fig. 1).

**Figure 1:**
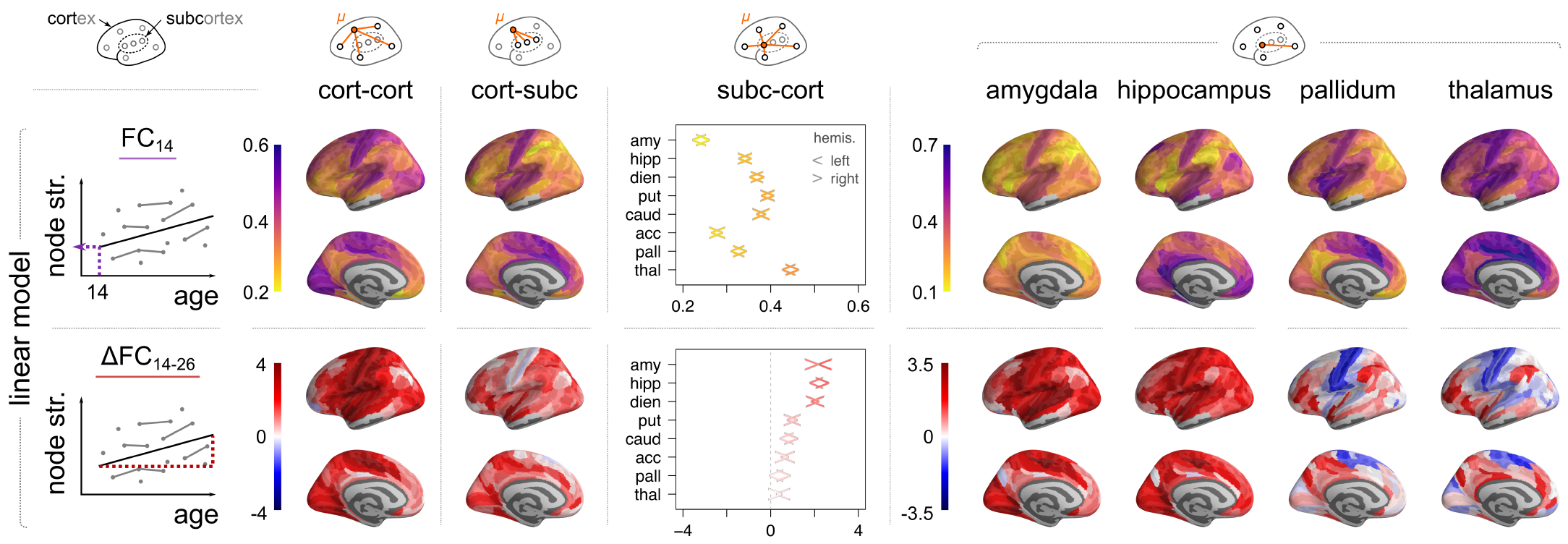
Regional strength of functional connectivity of cortical areas and subcortical nuclei at 14 years (*FC*_14_) and regional change in strength of connectivity during adolescence (∆*FC*_14−26_). Regional strength for each of 330 cortical and 16 subcortical nodes was regressed on a linear function of age for all participants (N=298; mixed effects model). Connectivity at age 14 was strongest for primary motor and sensory cortical areas, thalamus and striatal regions. Adolescent change in cortico-cortical connectivity strength was also strongly positive for primary motor and sensory areas; but these areas had reduced strength of connectivity to subcortical nuclei, i.e., ∆*FC*_14−26_ < 0. In contrast, connectivity between association cortex and subcortical nuclei, which was not strong at baseline, became more strongly positive during adolescence. In subcortico-cortical plots the left/right arrow corresponds to left/right hemisphere), and regions are ordered by average rate of change. Amygdala and hippocampus have weak cortical connectivity at baseline but the greatest rates of increase in cortical connectivity during adolescence. Thalamus has strong cortical connectivity at baseline but the greatest rates of decrease in primary cortical connectivity during adolescence. (For connectivity of the diencephalon, putamen, caudate and nucleus accumbens to cortex, see SI Fig. S5.)

At 14 years, all cortical regions had positive cortico-cortical connectivity strength and the most strongly connected nodes were located in primary motor and sensory cortical areas. Cortico-subcortical connectivity strength had a similar anatomical distribution, with stronger connectivity from primary cortical areas to subcortex, at baseline (Fig. 1). Age-related rates of change in connectivity strength were also regionally heterogeneous. Corticocortical connectivity strength increased in all regions during adolescence, most rapidly in primary motor and sensory cortical areas. However, age-related change in the strength of cortico-subcortical connectivity had a different anatomical distribution. The most positive rates of increase in connectivity to subcortical nodes were in association cortical areas, whereas some primary motor and sensory cortical areas had negative age-related changes in strength of connectivity to subcortical regions.

To investigate cortico-subcortical connectivity with more regional specificity, we estimated *FC*_14_ and ∆*FC*_14−26_ between each cortical area and each bilateral pair of 8 sub-cortical regions (Fig. 1 and SI Fig. S5). At baseline, the thalamus, the putamen and the pallidum were strongly connected to many cortical areas; whereas the amygdala and the accumbens had somewhat lower strength of cortical connectivity overall. The effect of age was also heterogeneous between subcortical regions. The amygdala, the hippocampus and the diencephalon had increased connectivity to many cortical areas; whereas the putamen, the pallidum, and the thalamus, had decreased strength of connectivity to primary somatomotor and premotor cortex, but increased strength of connectivity to frontal and parietal association cortex. While many of these age-related changes were statistically significant at an uncorrected probability threshold (*P* < 0.05), only the amygdala and the hippocampus had significantly increased connectivity to some cortical areas when controlling the false discovery rate (FDR < 0.05) (SI Table S3).

### Maturational index

Looking in more detail at the relationship between baseline FC and adolescent change in FC, we found that there was often a strong relationship between *FC*_14_ and ∆*FC*_14−26_ for the set of edges (correlations) connecting each regional node to the rest of the network. We defined the maturational index (MI) as the signed coefficient (Spearman’s *ρ*) of the linear relationship between edge-wise *FC*_14_ and ∆*FC*_14−26_ for each node (Fig. 2A) and we found that MI was often significantly non-zero by statistical tests including a spatial permutation test controlling for effects of spatial contiguity and hemispheric symmetry (*P*_spin_; SI Appendix). For example, the left somatosensory cortex had strongly positive MI, indicating that the edges with strongest FC at baseline showed the greatest positive increase in FC during adolescence. Conversely, left posterior cingulate cortex had strongly negative MI, indicating that the edges with weakest FC at baseline showed the greatest positive increase in FC during adolescence (Fig. 2B). To put it another way, in somatosensory cortex and other regions with MI > 0 there was a conservative mode of developmental change: connections that were already strong at 14 become stronger by the age of 26. Whereas, in posterior cingulate cortex and other regions with MI < 0, there was a disruptive mode of developmental change: connections that were weak at 14 got stronger by the age of 26 (or connections that were strong at baseline became weaker) (Fig. 2C,D).

**Figure 2:**
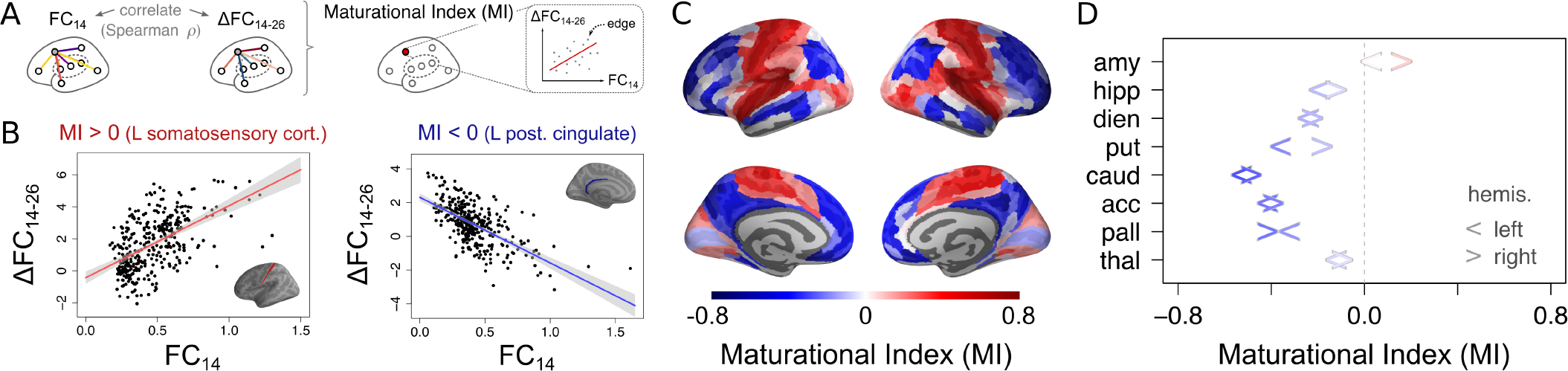
Maturational index. A) The maturational index (MI) for each brain region is defined as the correlation of edge-wise baseline *FC*_14_ versus rate of change ∆*FC*_14*−*26_. Panel B) illustrates this for two example regions, including a positive MI in left somatosensory cortex, and a negative MI in left posterior cingulate cortex. C) Visualisation of the Maturational Index for all cortical regions, and D) subcortical regions (the left/right arrow corresponds to left/right hemisphere).

Conservative changes in connectivity were concentrated in primary motor and sensory areas, corresponding to cytoarchitectonic classes 1 and 5 in the von Economo atlas (Vértes et al., 2016), and the insula (Fig. 3A). This anatomical distribution maps onto motor, ventral attention and visual networks previously defined by independent component analysis of adult resting state fMRI data (Fig. 3B) (Yeo et al., 2011). Disruptive changes in connectivity were concentrated in association cortex (von Economo class 2) and limbic cortex, corresponding to fronto-parietal, default mode and limbic resting state networks. Subcortical nodes were almost all characterized by disruptive development, with weak baseline connectivity to association cortex becoming stronger, or strong baseline connectivity to primary motor or sensory cortex becoming weaker (Fig. 2D).

**Figure 3:**
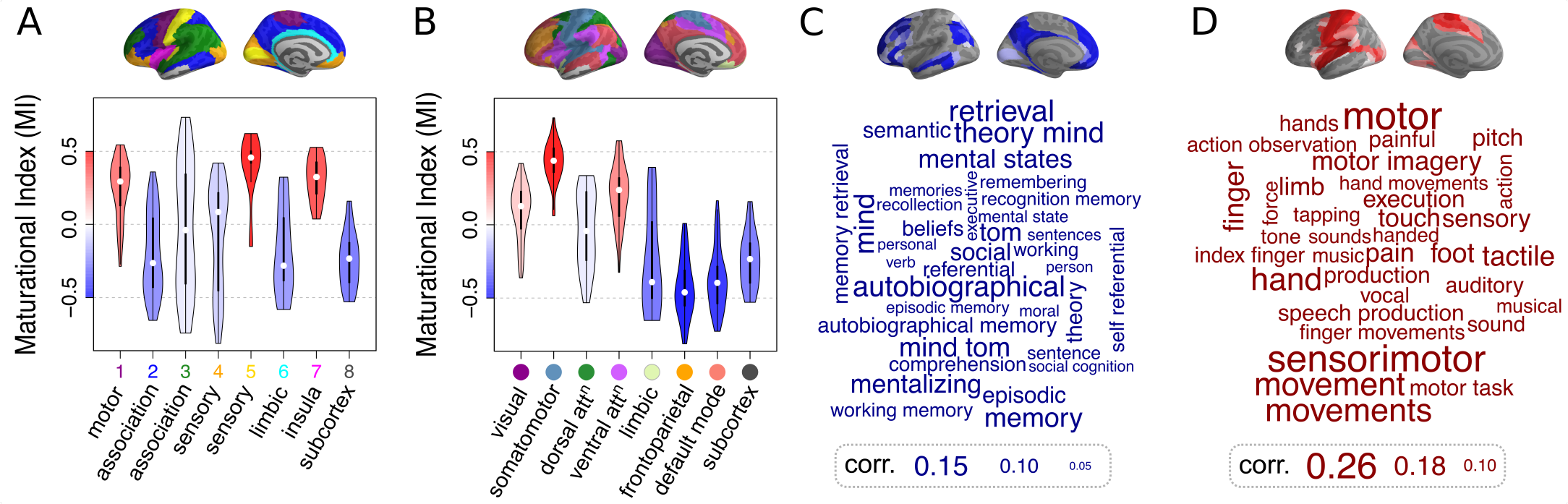
Maturational index in anatomical and psychological context. A) Distribution of maturational index for each cytoarchitectonic class of the von Economo atlas (Vértes et al., 2016), and B) for resting state networks derived from prior resting state FC analysis by Yeo (Yeo et al., 2011). In both cases, subcortical regions were considered as an additional eighth class/subnetwork. The violin plots are coloured by average MI within the corresponding class of regions. B) Word clouds of cognitive terms associated with cortical brain regions that have conservative (red) or disruptive (blue) modes of development (Neurosynth decoding; Yarkoni et al., 2011). The size of cognitive terms corresponds to the correlation of corresponding meta-analytic maps generated by Neurosynth with each of the two modes (top).

To ensure robustness of our results, we recomputed the MI using edge-wise *FC*_14_ and ∆*FC*_14−26_ derived from independent random half-splits of the data (2× 260 scans). The MI map averaged across 100 random half-splits was highly consistent with the map in Fig. 2 (SI Fig. S5).

### Contextualising adolescent change in FC

To assess the psychological relevance of these two modes of brain functional maturation, we used a meta-analytic tool (Neurosynth; Yarkoni et al., 2011) to identify cognitive processes that were specifically associated with disruptively vs conservatively developing cortical systems (Fig. 3C,D). We found that disruptive changes in FC were located in cortical areas that were specialised for memory, mentalising and social processing. Conversely, conservative changes in FC were located in cortical areas that were specialised for motor and sensory functions.

To assess the biological context of conservative and disruptive fMRI systems, we also compared the brain map of maturational index to several other relevant brain maps. For example, we estimated cortical thickness shrinkage at each cortical node in a cross-sectional dataset of structural MRI scans collected from 297 of the participants in this fMRI study (Whitaker et al., 2016). We found that cortical areas with the most negative rates of thickness change during adolescence (or fastest shrinkage) had the most negative MI (Spearman’s *ρ* = 0.16, *P* = 0.0033, *P*_spin_ = 0.026; Fig. 4A). This indicates that disruptive adolescent changes in FC are co-located with well-replicated adolescent changes in cortical structure. However, other structural MRI parameters were not significantly co-located with MI in this sample (SI Fig. S7).

**Figure 4:**
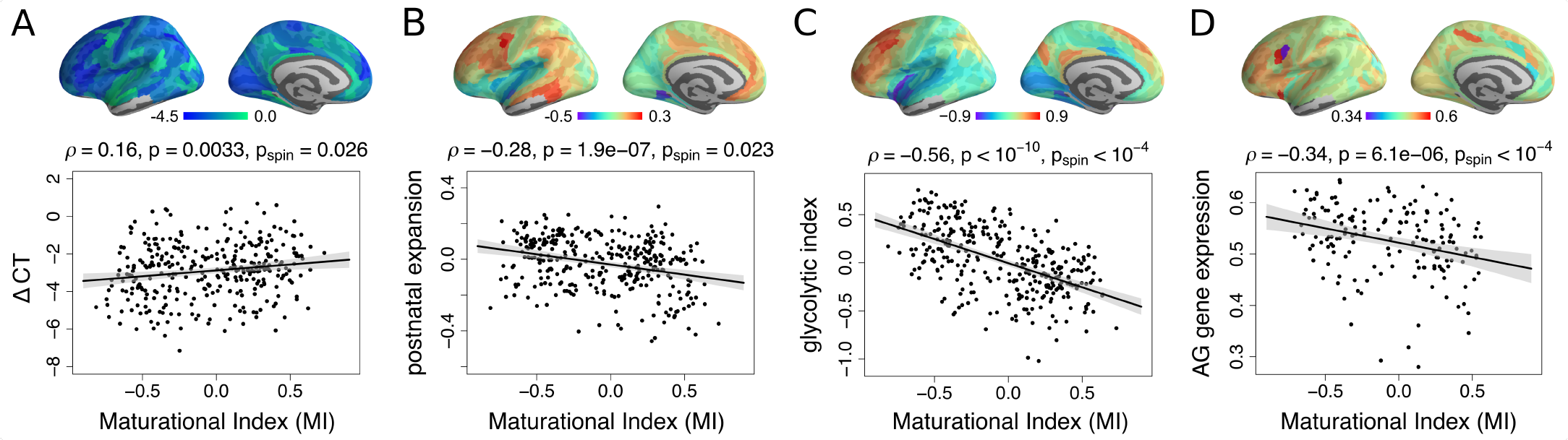
Disruptive and conservative modes of fMRI maturation in developmental and metabolic context. A) Maturational index was positively correlated with ∆*CT* (Whitaker et al., 2016) - regions which had disruptive development (MI<0) had faster rates of cortical thickness (CT) shrinkage during adolescence. B) MI was negatively correlated with a prior map of postnatal cortical surface area (Vaishnavi et al., 2010) - disruptive maturation was greater in regions that showed greatest expansion after birth. C) MI was negatively correlated with a prior map of the glycolytic index, a measure of aerobic glycolysis (AG; Vaishnavi et al., 2010); and D) MI was negatively correlated with a prior map of brain regional expression of AG-related genes (Hawrylycz et al., 2012; Arnatkeviciute et al., 2019).

Additionally, we compared the maturational index map (Fig. 2C) to nine independently produced maps of a range of brain functional and developmental parameters, including: (i) metabolic rates of glucose, oxygen and aerobic glycolysis (AG) measured by PET (Vaishnavi et al., 2010); (ii) microarray measures of gene expression for 116 genes previously associated with AG (Goyal et al., 2014) and extracted from the Allen Human Brain Atlas (Hawrylycz et al., 2012) as in Arnatkeviciute et al. (2019); (iii) evolutionary and post-natal surface expansion of the cortex (Hill et al., 2010); and (iv) areal scaling of the cortical surface (Reardon et al., 2018).

We found that disruptive cortical regions (with negative MI) had higher metabolic rates of glucose (Spearman’s *ρ* = −0.41, *P* < 10^−10^, *P*_spin_ = 0.0013), higher rates of AG as measured by the glycolytic index (Spearman’s *ρ* = −0.56, *P* < 10^−10^, *P*_spin_ < 10^−4^), higher expression of AG-related genes (Spearman’s *ρ* = −0.34, *P* < 6.1·10^−6^, *P*_spin_ < 10^−4^), and faster rates of postnatal surface expansion (Spearman’s *ρ* = −0.28, *P* < 1.9⋅10^−7^, *P*_spin_ =0.023). For further details see Fig. 4 and SI Fig. S7.

### Head motion and distance-related change in connectivity

We found no evidence of an age-related change in participants’ overall in-scanner motion, quantified using framewise displacement (FD) (*t* = 1.36, *df* = 226, *P* = 0.18) (Power et al., 2012). However, there was a positive average correlation between edge-wise FC and head movement (*r* = 0.25) which was corrected by regressing average participant motion (mean FD) from FC at each edge (SI Fig. S2). After these pre-processing steps, there was no evidence that the Euclidean distance between regional nodes was related to age-related changes in FC (Spearman’s *ρ* < −0.01, P = 0.95), indicating that adolescent maturation of FC was not greater for longer (or shorter) distance connections.

## Discussion

We have reported results from an accelerated longitudinal study of adolescent development of functional connectivity (FC) in the healthy human brain. In a large, population-representative sample of fMRI data, balanced for age and sex, and carefully controlled for head motion, we found evidence for two modes of maturational change in the age range 14 to 26 years, which we called conservative and disruptive.

The conservative mode of change was consolidating, or making stronger over the course of adolescence, the FC of regional nodes that were already highly connected hubs of the resting state fMRI network at the start of adolescence. Conservatively, “the rich get richer”. Conservative maturation was concentrated in motor, primary sensory and insula areas of cortex defined cytoarchitectonically; corresponding to visual, somatomotor and ventral attentional networks defined by prior analysis of adult fMRI data; and specialised for motor, somatosensory and auditory functions, by meta-analysis of prior task-related fMRI data.

The disruptive mode of change was to make FC stronger in cortical nodes where it was relatively weak at age 14, and to make it weaker in subcortical nodes where it was relatively strong at the start of adolescence. Disruptively, “the rich get poorer and the poor get richer”. Disruptive maturation was characteristic of association and limbic cortex, corresponding to default mode, fronto-parietal and limbic fMRI networks, and specialised for memory, theory of mind, and social cognition.

Some convergent fMRI connectivity results have been recently reported. Gu et al. (2015) studied 780 participants aged 8-22 years (each scanned once) and measured FC within and between different cognitive systems. They found that motor and sensory systems showed developmental increases in within-system connectivity along with decreased between-system connectivity, thus reinforcing the connectivity profile established earlier on in childhood.

We hypothesised that the disruptive pattern of changes in FC was consistent with the concept that synaptic connectivity of association cortical and subcortical brain systems was developmentally re-modeled during adolescence to enable the emergence of adult social and executive skills and self-awareness. We explored various aspects of this hypothesis by comparing the anatomical map of the maturational index (MI), defining disruptively (or conservatively) developing brain systems, to prior brain maps of structural, genomic and metabolic parameters of adolescent development.

For example, positron emission tomography (PET) has been used to map the fraction of glucose that is oxidized to supply cells with energy in the form of ATP, versus the fraction undergoing aerobic glycolysis (AG), a non-oxidative metabolic pathway in the presence of oxygen. Developmental PET studies have shown that association cortical regions display significant AG throughout adolescence and into adulthood (Vaishnavi et al., 2010; Goyal et al., 2014), while primary sensory and motor regions progressively reduce glucose metabolism by AG (and increase oxidative metabolism) from early childhood into adolescence (Vaishnavi et al., 2010; Goyal et al., 2014). The maintenance of high aerobic glycolysis in adolescent and adult association cortex could indicate delayed or persistent development of those regions, requiring metabolic energy provided specifically by AG in support of processes such as synaptic remodeling (Goyal et al., 2014).

We found that the prior PET map of glycolytic index (GI) was highly correlated with the fMRI map of maturational index (MI). Association cortical and subcortical regions with MI < 0 had GI > 0; whereas motor and sensory cortical areas with MI > 0 had GI < 0. This result was corroborated by the significant spatial correlation between a prior map of expression of AG-related genes (coding for 116 proteins) and the fMRI map of MI. Disruptively developing brain regions had higher levels of AG-related genes than conservatively developing regions. We regard these convergent results as indicating that disruptive adolescent development of fMRI connectivity represents a metabolically expensive process of synaptic re-modelling in association cortex and subcortex.

We also found significant anatomical correspondence between the fMRI map of MI and the map of cortical shrinkage derived from structural MRI data in the same sample. Cortical shrinkage is the most well-replicated result in MRI studies of adolescent brain development and has been mechanistically explained as a marker of synaptic pruning and/or intra-cortical myelination. Association cortical areas have both faster rates of shrinkage and more disruptive changes to their FC during adolescence. Another structural measure of developmental activity was provided by a prior map of post-natal expansion of cortical surface area (Hill et al., 2010). Association cortex has both greater surface area expansion and more disruptive development of FC (Fig. 4). We regard these results as convergently indicating that disruption of FC between regions is co-located with cortical systems that are most structurally active in their development during adolescence.

Finally, we considered the psychological implications of these two modes of adolescent brain development. Conservative systems were specialised for relatively basic sensory and motor functions that would normally have been operational since early childhood. Disruptive systems were specialised for a range of “higher-order” functions, such as working memory, theory-of-mind and autobiographical memory, which are known to be relatively late-maturing social and cognitive processes and characteristic of psychological development of adolescence. We can hypothesise that the neurodevelopmental programme represented by disrupted FC drives the emergence of more sophisticated socialising, mentalizing and executive skills as young people grow into independent adults. To test this hypothesis more rigorously will require design and analysis of longitudinal studies of development of cognitive functions as well as fMRI connectivity. However, if indeed normal emergence of adult emotional and social cognitive faculties is dependent on disruptive maturation of association cortical and subcortical FC, then it is also predictable that incidence of psychiatric disorders or subclinical psychopathology in adolescence could arise from abnormal trajectories of disruptive maturation. We have found that trajectories of intra-cortical myelination in fronto-striatal systems are related to individual differences in compulsivity and impulsivity (Ziegler et al., 2018). It is also noteworthy that both the default mode network (Wise et al., 2017) and cortico-subcortical connectivity (Heller, 2016) have been previously implicated in the pathophysiology of major depressive disorder.

### Methodological issues

Strengths of the study include the accelerated longitudinal design and the balanced sample of healthy young people stratified by age and sex. Informed by prior insights (Power et al., 2012; Satterthwaite et al., 2012, 2013), we prospectively used a multi-echo fMRI sequence to disambiguate BOLD components from other, non-neuronal sources of fMRI dynamics, such as head movement. We carefully quality controlled all fMRI data to ensure that there were no discernible biases in estimation of the correlation between regional mean time series that were attributable to head motion. The sample size was evidently sufficient to detect some major developmental changes in FC but a larger sample would be necessary to investigate more powerfully the links between structural and functional MRI markers of brain development. The gene expression maps used for the co-location analysis of AG-related genes were derived from post-mortem data on 6 adult brains.

## Conclusion

Our results indicate substantial reorganisation of association cortical and sub-cortical connectivity in the adolescent functional connectome (Váša et al., 2017). We propose that these macroscopic, maturational changes reflect microscopic synaptic remodelling of large-scale brain circuits that are relevant to the healthy emergence of adult faculties in adolescence.

## Methods

### Participants

A demographically balanced cohort of 306 healthy adolescents (153 females) aged 14-26 years, scanned a total of 556 times, was considered for inclusion in this study. There were approximately equal numbers of male and female participants (~30) in each of 5 age-defined strata at baseline: 14-15 years inclusive, 16-17 years, 18-19 years, 20-21 years and 22-24 years. The study was ethically approved by the National Research Ethics Service and was conducted in accordance with NHS research governance standards.

### MRI acquisition and pre-processing

Scanning took place at three sites, all operating identical 3T MRI systems (Magnetom TIM Trio, Siemens Healthcare, VB17 software version). Resting-state fMRI data were acquired using a multi-echo echoplanar imaging (ME-EPI) sequence (Barth et al., 1999): TR = 2.42 s; GRAPPA with acceleration factor = 2; matrix size= 64 × 64 × 34; FOV = 240 × 240 mm; in-plane resolution = 3.75 × 3.75 mm; slice thickness = 3.75 mm with 10% gap, sequential slice acquisition, 34 oblique slices; bandwidth = 2368 Hz/pixel; echo times (TE) = 13, 30.55 and 48.1 ms.

For pre-processing of functional scans, we used multi-echo independent component analysis (ME-ICA; Kundu et al., 2012, 2013) to identify the sources of variance in the fMRI time series that scaled linearly with TE and could therefore be confidently regarded as BOLD signal. Other sources of fMRI variance, such as head movement, which were not BOLD-related and therefore did not scale with TE, were identified by ME-ICA and discarded. The retained independent components, representing BOLD contrast, were optimally recomposed to generate a broadband denoised fMRI time series at each voxel (Posse et al., 1999). This was bandpass filtered by the discrete wavelet transform (Daubechies 4 wavelet), resulting in a BOLD signal oscillating in the frequency range 0.025-0.111 Hz (wavelet scales 2 and 3). Geometric re-alignment of scans was used to estimate 6 motion parameters for each participant (3 translation and 3 rotation parameters). These were used to calculate an overall estimate of motion - the framewise displacement (FD; as in Power et al. (2012)). For each participant, mean FD was calculated by averaging the FD time series. A total of 36 scans were excluded due to high in-scanner motion (*µ*(FD) > 0.3 mm or *max* (FD) > 1.3 mm) and other quality control criteria (see SI).

### Parcellation and FC estimation

Functional MR images were parcellated using prior anatomical templates. For principal analyses, we used a multi-modal parcellation of cortex into 360 bilaterally symmetric regions (Glasser et al., 2016), as well as 16 subcortical regions from FreeSurfer software (Filipek et al., 1994), yielding a total of 376 regions. Regional BOLD signals were estimated by averaging pre-processed time series over all voxels in each parcel. Some regions (particularly near frontal and temporal poles) were excluded because of low regional mean signal, defined by a low Z-score of mean signal intensity in at least one subject (Z < −1.96); this resulted in the exclusion of 30 cortical regions. To test the sensitivity of our results to the cortical parcellation, we repeated analyses using a sub-parcellation of the Desikan-Kiliany anatomical atlas (Desikan et al., 2006) into cortical 308 parcels of approximately equal surface area (~5cm^2^; Romero-Garcia et al., 2012) (see SI for further details).

After pre-processing and quality control, regional fMRI time series were available at 330 cortical areas and 16 subcortical structures for 298 participants (151 females), scanned a total of 520 times.

Functional connectivity matrices were estimated for each scan by compiling all possible pair-wise inter-regional Pearson’s correlations between regional time series. Agerelated change in FC was modelled using linear mixed effect models that included age as the main fixed effect of interest, sex and scanner-site as fixed effect covariates, as well as a subject-specific intercept as a random effect (see SI for further details).

## Data and code

Processed data and code to reproduce results will be made available upon publication of the manuscript.

## Author contributions

P.F., R.J.D., P.B.J., I.M.G. and E.T.B. designed the NSPN study; F.V, P.E.V., and E.T.B. designed analyses; F.V., R.R.G., M.G.K., J.S., K.J.W., and M.M.V. processed and quality controlled data; J.S., P.K., and A.X.P. contributed new analytic tools; F.V. and P.E.V. analyzed data; F.V., P.E.V., and E.T.B. wrote the paper.

## Acknowledgments

This study was supported by the Neuroscience in Psychiatry Network, a strategic award by the Wellcome Trust (095844/Z/11/Z). Additional support was provided by the National Institute for Health Research (NIHR) Cambridge Biomedical Research Centre and the Medical Research Council (MRC)/Wellcome Trust Behavioural and Clinical Neuroscience Institute. FV was supported by the Gates Cambridge Trust. PEV was supported by the MRC (MR/K020706/1), is a Fellow of MQ: Transforming Mental Health (MQF17/24), and is a Fellow of the Alan Turing Institute funded under EPSRC (EP/N510129/1). PF is a NIHR Senior Investigator (NF-SI-0514-10157), and was in part supported by the NIHR Collaboration for Leadership in Applied Health Research and Care (CLAHRC) North Thames at Barts Health NHS Trust. ETB is a NIHR Senior Investigator. The views expressed are those of the authors and not necessarily those of the NHS, the NIHR or the Department of Health and Social Care.

