## Supplementary material for "Conservative and disruptive modes of adolescent change in brain functional connectivity": SI

### Supplementary Methods

*Participants*

A demographically balanced cohort of 306 adolescents (153 females) aged 14-26 years was considered for inclusion in this study. Of these, 313 were scanned at "baseline", with approximately 60 participants in each of 5 age-defined strata: 14-15 years inclusive, 16-17 years, 18-19 years, 20-21 years and 22-26 years. Subsequently, 28 participants were scanned approximately 6 months later, and 215 participants were scanned approximately a year later. Thus, a total of 556 scans were acquired. Following exclusion of certain scans, 520 scans were retained; see below and Fig. S1.

Participants provided informed written consent for each aspect of the study, and parental consent was obtained for those aged 14–15 years. None of the included participants were currently being treated for a psychiatric disorder or for drug or alcohol dependence; had a current or past history of neurological disorders or trauma; or had a learning disability. The study was ethically approved by the National Research Ethics Service and was conducted in accordance with NHS research governance standards.

*MRI acquisition and pre-processing*

Scanning took place at three sites, all operating identical 3T MRI systems (Magnetom TIM Trio, Siemens Healthcare, VB17 software version) with standard 32-channel radio-frequency (RF) receive head coil and RF body coil for transmission. Resting-state fMRI data were acquired using a multi-echo echoplanar imaging (ME-EPI) sequence with online reconstruction (Barth et al., 1999): repetition time (TR) = 2.42 s; GRAPPA with acceleration factor = 2; flip angle = 90°; matrix size = 64 x 64 x 34; FOV = 240 x 240 mm; in-plane resolution = 3.75 x 3.75 mm; slice thickness = 3.75 mm with 10% gap, sequential slice acquisition, 34 oblique slices; bandwidth = 2,368 Hz/pixel; echo time (TE) = 13, 30.55 and 48.1 ms. Following processing of individual structural scans using FreeSurfer v5.3.0 (including skull-stripping, segmentation of cortical grey and white matter and reconstruction of the cortical surface and grey-white matter boundary) (Fischl et al., 1999), all scans were stringently quality controlled by re-running the reconstruction algorithm after the addition of control points and white matter edits (as described previously; Whitaker et al., 2016; Váša et al., 2017).

Coregistration of structural and functional scans was visually assessed. For pre-processing of functional scans, we used multi-echo independent component analysis (ME-ICA; Kundu et al., 2012, 2013) to identify the sources of variance in the fMRI time series that scaled linearly with TE and could therefore be confidently regarded as indicative of BOLD contrast. Other sources of fMRI variance, such as head movement, which were not BOLD-dependent and therefore did not scale with TE, were identified by ME-ICA and discarded. The retained independent components, representing BOLD contrast, were optimally recomposed to generate a broadband denoised fMRI time series at each voxel (Posse et al., 1999). This was bandpass filtered by the discrete wavelet transform (Daubechies 4 wavelet), resulting in a BOLD signal oscillating in the frequency range 0.025-0.111 Hz (wavelet scales 2 and 3).

During processing, re-alignment of scans was used to estimate 6 motion parameters for each participant (3 translation parameters and 3 rotation parameters). Subsequently, these were used to calculate an overall estimate of motion - the framewise displacement (FD), defined as the sum of the absolute derivatives of the six motion parameters, following conversion of rotational parameters to distances by computing the arc length displacement on the surface of a sphere with radius 50 mm (as in Power et al. (2012) and Patel et al. (2014)):

$$FD_t = \sum_d \left| d_{(t-1)} - d_t \right| + 50 \cdot \frac{\pi}{180} \cdot \sum_r \left| r_{(t-1)} - r_t \right| \quad (1)$$

where $d$ denotes translation distances $\{x,y,z\}$, and $r$ denotes rotation angles $\{\alpha,\beta,\gamma\}$. For each participant, mean FD was calculated by averaging the FD time series.

*Parcellation and network construction*

Each subject's cortex was parcellated using two parcellations: a recent multi-modal parcellation into 360 bilaterally symmetric regions based on data from the Human Connectome Project (Glasser et al., 2016), as well as a sub-parcellation of the Desikan-Kiliany anatomical atlas (Desikan et al., 2006) into 308 parcels of approximately equal surface area ($\sim$5cm$^2$; Romero-Garcia et al., 2012). Additionally, 16 subcortical regions were provided by FreeSurfer software (Filipek et al., 1994), and included bilateral pairs of the following regions: thalamus, caudate, putamen, pallidum, hippocampus, amygdala, nucleus accumbens and the ventral diencephalon. Thus, the combination of each of the two cortical parcellations with the subcortical regions yielded parcellations of cortex and subcortex into 376 regions (henceforth referred to as the HCP parcellation) and 324 regions (henceforth referred to as the DK-sub parcellation) respectively.

Regional BOLD time series were estimated by averaging time series over all voxels in each parcel. Some regions (particularly near the frontal and temporal poles) were excluded because of low regional mean signal, defined by a low

Z-score of mean signal intensity in at least one subject (Z $<$ -1.96); this resulted in the exclusion of 30 cortical regions for the HCP parcellation, and 26 regions for the DK-sub parcellation, primarily in inferior temporal and pre-frontal cortex. Thus, the final number of regions (both cortical and subcortical) used for analyses was 346 in the HCP parcellation, and 298 in the DK-sub parcellation.

Finally, FC matrices were constructed by cross-correlating the BOLD time series between pairs of the retained regions (using Pearson's correlation).

*Exclusion criteria and final sample*

A total of 36 scans were excluded. Of these: 17 scans were excluded due to high in-scanner motion (defined as mean FD $>$ 0.3 mm or maximum FD $>$ 1.3 mm), 9 due to coregistration errors, 7 due to a lack of convergence of the ME-ICA algorithm, 2 due to parcellation errors, 1 due to extensive dropout (defined above).

Following quality control and participant exclusion, 298 adolescents (151 females), scanned a total of 520 times, were included in the study. Of these, 298 scans were acquired at "baseline", 27 were acquired approximately 6 months later, and 195 were acquired approximately a year later (Fig. S1). Within the final included sample, 95 participants were scanned once, 184 participants were scanned twice, and 15 participants were scanned three times. Thus, the study included 520 unique FC matrices capturing pair-wise FC of 346 (HCP) and 298 (DK-sub) cortical and subcortical regions.

*Effects of motion on functional connectivity*

We first verified whether participants' overall motion (quantified using time-averaged FD) changes as a function of age (using linear mixed effects models; see details below). Moreover, to evaluate potential residual effects of motion on FC, we calculated the correlation between FC at each edge and participants' motion (across participants). Subsequently, to assess whether motion affected FC in a distance-dependent manner, we correlated the upper triangular part of this matrix (of correlations between FC and motion) to the upper triangular part of a matrix of Euclidean distances between the centroids of all regions, as in Satterthwaite et al. (2012) (Fig. S2).

*Maturation of functional connectivity*

Maturation of FC as a function of age was modelled using linear mixed effect models. These models included age as the main fixed effect of interest, as well as sex and scanner site as fixed effect covariates, as well as a subject-specific intercept as a random effect, as follows:

$$FC = \beta_{age} \cdot age + \beta_{sex} \cdot sex + \beta_{site} \cdot site + \gamma_{subj} \cdot (1|subj) + \epsilon \tag{2}$$

where $FC$ refers to functional connectivity (at one of several possible levels of spatial resolution; see below); $\beta$ refers to coefficients for the fixed effects, $\gamma$ refers to coefficients for random effects and $\epsilon$ represents the residual error. We further explored inclusion of an interaction term between age and sex, as well as intra-cranial volume (ICV) as additional covariates. These models presented poorer fits (as indicated by increased AIC and BIC) than models without these terms fitted to the same data. Therefore, we focused on fixed effects of age and sex only.

Additionally, we fitted locally adaptive mixed effect smoothing splines (generalized additive mixed models) to regional trajectories, better suited than quadratic models at modelling non-linear trajectories (Fjell et al., 2010). Specifically, we used the *gamm* function in R, with effect of age modelled as a weighted sum of 10 cubic b-splines with knots placed at quantiles of the data and smoothing optimized using restricted maximum likelihood (REML) Reiss et al. (2014). We inspected for evidence of non-linearity using the degrees of freedom ($df$) of the smoothing splines. However, we found no substantial evidence of non-linearities, and thus focused on linear trajectories (see Supplementary results below, and Fig. S4).

Linear mixed effect models were fitted at multiple spatial scales: (i) globally, at the level of the correlation distribution, (ii) regionally, using average nodal FC and (iii) edge-wise.

*Global maturation of functional connectivity*

To evaluate maturational shifts in the functional correlation distribution, we first summarised each individual distribution using its four moments: the mean, variance, skewness and kurtosis, fitting linear models as a function of age to each. Further, to more precisely characterise potential local age-shifts in the correlation distribution, we fitted linear models to the value of correlation at each percentile of the distribution, and compared the effect sizes across percentiles.

*Nodal maturation of functional connectivity*

We next focused on individual regions using node strength, defined as the average FC of a node to other nodes. In addition to this overall estimate of node strength, we recalculated node strength over subsets of cortical edges (between the 330 cortical regions and all 346 regions) and subcortical edges (between the 16 subcortical regions and all 346 regions). For all both nodal strength maps (cortical-all and subcortical-all), we characterised regional development of FC by fitting linear models of changes in node strength as a function of age. In addition to the effect size of changes as a function of age ($\Delta FC_{14-26}$), we extracted predicted FC at age 14 years ($FC_{14}$). Moreover, we constructed maps of subcortico-cortical maturation of FC for each of the eight bilateral pairs of subcortical regions listed above. Maps of the development of FC to each cortical node were obtained by fitting linear models to the average of the two edges between each bilateral pair of subcortical regions and each cortical region.

*Edge-wise maturation of functional connectivity*

Finally, the most fine-grained description of FC development was obtained by fitting linear models as a function of age to each edge (Pearson's correlation) in the FC matrix. This led to matrices of the edge-wise effect sizes of age ($\Delta FC_{14-26}$), as well as edge-wise FC at 14 ($FC_{14}$). These matrices were then explored in several ways. First, we inspected potential effects of the distance of functional connections on their maturation, by evaluating Spearman's correlation between the upper triangular parts of the matrices of Euclidean distance and effects of age on FC (Fig. S8). Subsequently, we evaluated at each node the relationships (Spearman's $\rho$) between corresponding edge-wise parameters: $FC_{14}$ and $\Delta FC_{14-26}$, which we term "Maturational Index" (MI) in the main text.

*Spatial permutation "spin" test for comparison of cortical maps*

When relating pairs of regional cortical maps to each other across regions, non-independence among regions (due to spatial auto-correlation amongst neighbouring parcels) should be taken into account. Here, we used a regional spherical permutation test as implemented in (Váša et al., 2017), following earlier implementation of such tests at the vertex level (Alexander-Bloch et al., 2013; Vandekar et al., 2015).

The spatial permutation test was implemented in the following manner: First, we obtained the coordinates of each of our 330 regions on the FreeSurfer spherical projection of the parcellation. We next rotated these coordinates about the three axes (x: left-right, y: rostral-caudal, z: dorsal-ventral) at three randomly generated angles, $\theta_x$, $\theta_y$ and $\theta_z \in [0,2\pi)$, using the following rotation matrices:

$$R_x(\theta) = \begin{bmatrix} 1 & 0 & 0 \\ 0 & cos(\theta) & -sin(\theta) \\ 0 & sin(\theta) & cos(\theta) \end{bmatrix} \tag{3}$$

$$R_y(\theta) = \begin{bmatrix} cos(\theta) & 0 & sin(\theta) \\ 0 & 1 & 0 \\ -sin(\theta) & 0 & cos(\theta) \end{bmatrix} \tag{4}$$

$$R_z(\theta) = \begin{bmatrix} cos(\theta) & -sin(\theta) & 0 \\ sin(\theta) & cos(\theta) & 0 \\ 0 & 0 & 1 \end{bmatrix} \tag{5}$$

Since each hemisphere is projected onto the sphere separately, the rotation was applied to both hemispheres. To preserve hemispheric symmetry, the same random angles were applied to both hemispheres, with the caveat that the sign of the angles was flipped for the rotations around the y and z axes; i.e.:, $\theta_{yR}$ = -$\theta_{yL}$ and $\theta_{zR}$ = -$\theta_{zL}$ (but $\theta_{xR} = \theta_{xL}$).

Following rotation of the sphere, coordinates of the rotated regions were matched to coordinates of the initial regions, using Euclidean distance and proceeding in a descending order of average Euclidean distance between pairs of regions on the rotated and unrotated spheres (i.e.: starting with the rotated region that is furthest away, on average, from the unrotated regions). The matching then provides a mapping from the set of regions to itself, that allows any regional measure to be permuted while controlling for spatial contiguity and hemispheric symmetry.

P-values for the correlation between two maps were obtained by comparing the empirical value of Spearman's $\rho$ to a null distribution of 10'000 Spearman correlations, between one empirical map and a set of 10'000 spatially permuted versions of the other map. Each analysis correlating values from two cortical maps is reported with both the P-value corresponding to the Spearman correlation (P), as well as a P-value derived from the spherical permutation ($P_{\text{spin}}$).

*Contextualising findings*

*von Economo and Yeo atlases*

We mapped the von Economo atlas of 7 cytoarchitectonic classes (von Economo and Koskinas, 1925) to our cortical parcellations. For our in-house sub-parcellation of the Desikan-Killiany atlas (Romero-Garcia et al., 2012) into 308 cortical regions, we used a manual labeling of Desikan-Killiany regions to von Economo classes, previously described and used in (Vértes et al., 2016). For the HCP parcellation (Glasser et al., 2016), we first computed overlap between each HCP parcel and each cytoarchitectonic class on the surface (using CIVET software; Ad-Dab'bagh et al., 2006), before assigning each parcel to the class that it overlapped most. Due to symmetry of our original von Economo atlas, we computed overlap across both hemispheres simultaneously, resulting in a symmetric assignment of von Economo classes to HCP parcels. In addition to the seven classes used in (Vértes et al., 2016; Váša et al., 2017), we considered an additional "class", consisting of the eight bilateral pairs of subcortical regions included in our anatomical atlas. The Yeo atlas (Yeo et al., 2011) was mapped in a similar manner, except that the original vertex-wise atlas was used.

*Neurosynth*

To perform the Neurosynth (Yarkoni et al., 2011) functional meta-analysis, surface maps were converted from Freesurfer *fsaverage* to MNI space using barycentric interpolation via the SurfToSurf command in AFNI (Cox, 1996). The resultant cortical surface maps were then uploaded to Neurovault and analyzed with the Neurosynth "decoder" function.

*Cortical maps*

The PET maps (remapped from (Vaishnavi et al., 2010; Glasser et al., 2014; Reardon et al., 2018)) were provided via personal correspondence with the research groups that had generated the data. We first converted these maps to the *fsaverage6* surface template using standard transformations provided in the HCP Connectome Workbench (v1.2.2) software package (https://www.humanconnectome.org/software/connectome-workbench). Next, we transformed the *fsaverage6*-aligned map to the MNI surface space by a direct surface-to-surface transformation in which values at vertices in the target coordinate matrix were assigned by ones of their nearest neighbor vertices within a 10 mm radius in the source coordinate matrix. To relate these vertex-level maps to our parcel-based results, we first averaged the vertex values within each parcel. Due to the availability of only the group-level PET maps, the glycolytic index (GI) was performed at the vertex level by taking the scaled (by 1000) residuals of the group average PET-CMRGlu map regressed on the group average PET-CMRO2 map as performed at the subject-level in (Vaishnavi et al., 2010).

*Gene expression analyses*

Prior work by Goyal et al. has identified 116 genes whose expression pattern across the cortex was correlated with the extent to which different brain regions rely on aerobic glycolysis (Goyal et al., 2014). Here, we used data from the Allen Human Brain Altas (AHBA) – a transcriptomic dataset created by the Allen Institute for Brain Sciences (http://human.brain-map.org) (Hawrylycz et al., 2012) – to ask whether the mean expression value of these 116 genes also correlates with the maturational index derived from fMRI. The AHBA dataset is based on six donor brains from three Caucasian, two African-American and one Hispanic donors. Their ages were 57, 55, 49, 39, 31 and 24 years. For the results reported in the main manuscript we used the data made available alongside the recent paper (Arnatkeviciute et al., 2019) which had undergone the following processing steps (as described in the paper): "(i) confirming and updating probe-to-gene annotations using the latest available data; (ii) background filtering, where expression values that do not exceed background are removed; (iii) probe selection, which, for genes indexed by multiple probes, involves selecting a single representative measure to represent the expression of that gene; (iv) sample assignment, where tissue samples from the AHBA are mapped to specific brain regions in an imaging dataset (in this case mapping to the parcels in the HCP parcellations); (v) normalization of expression measures to account for inter-individual differences and outlying values".

We then selected the genes from Goyal et al. (2014), averaged their expression across cortical brain regions and correlated this vector with the maturational index. Note that four of the six AHBA donor brains were only sampled in the left hemisphere, due to the presumed similarity of gene expression between hemispheres. For this reason we only use brain regions from the left hemisphere in main text Figure 4D.

In the supplement, we show that the results are unchanged when using a previously published form of the AHBA data preprocessed and matched to the DK sub-parcellation as described in (Romero-Garcia et al., 2018) (including slightly different criteria for each of the preprocessing steps described above).

### Supplementary results

*Effects of motion on functional connectivity*

We found that the average participant motion (as measured by FD) did not change with age. However we did find residual signatures of motion in our data, including relationships between average participant motion and FC at several spatial scales, and a weak distance-dependence of the relationship between FC and motion at each edge. In the present work, we have removed these residual effects using regression, as has been applied previously (Gu et al., 2015) (Fig. S2).

*Global maturation of functional connectivity*

The correlation between fMRI time series for a pair of cortical and/or subcortical regions was generally positive indicating FC through coherent in-phase low frequency BOLD oscillations. The global parameters of the correlation distributions, estimated over all 70,500 edges in each correlation matrix, shifted over adolescence. Between 14 and 26 years, there was a weak but significant trend for the average or mean correlation to increase ($t(221) = 2.1$, $P = 0.035$). There were also stronger age-related effects on the higher-order parameters of the correlation distribution: variance or spread increased ($t(221) = 5.2$, $P = 3.8 \cdot 10^{-7}$)); skewness decreased ($t(221) = -2.6$, $P = 0.0097$)); and kurtosis or peakiness decreased ($t(221) = -4.9$, $P = 2.1 \cdot 10^{-6}$). We also characterised changes in the correlation distribution more precisely by fitting linear models to estimate the effect of age on the value of correlation at each percentile. The edges that were most strongly connected at baseline (14 years) demonstrated the most rapid rates of further increase in positive correlation over the course of adolescence (Fig. S3).

*Nodal maturation of cortico-subcortical functional connectivity*

To investigate cortico-subcortical connectivity with more regional specificity, we estimated $FC_{14}$ and $\Delta FC_{14-26}$ between each cortical area and each of 8 subcortical regions (Fig. S5). While many of these age-related changes were statistically significant at an uncorrected probability threshold ($P < 0.05$), only the amygdala and the hippocampus had significantly increased connectivity to some cortical areas when controlling the false discovery rate (FDR $< 0.05$) (Table S3).

*Fitting of smoothing splines to edge-wise FC trajectories*

To inspect maturational trajectories for the potential presence of non-linearities, we fitted locally adaptive mixed effect smoothing splines to edge-wise data. Subsequently, we extracted the effective degrees of freedom ($df$) of the smoothing splines ($df = 2$ corresponds to a linear trajectory).

Most trajectories were linear: of all edges (in the HCP parcellation), 71.7% trajectories had $df < 2.1$, and 80.0% had $df < 2.5$ (Fig. S4A). The cortical distribution of average nodal $df$ (averaged across all of a node's edges) showed that regions with $df > 2.5$ were scattered across cortex, and not systematically located within specific anatomical or functional areas (Fig. S4B). Finally, the maturational index (MI) showed a weak relationship with average nodal $df$ (Spearman's $\rho = -0.097$, $P = 0.07$, $P_{\text{spin}} = 0.027$), with a slight tendency of "disruptive" regions (with MI $< 0$) to show more (weakly) non-linear trajectories (Fig. S4C). This is in line with prior work showing adolescent non-linearities in association-cortical trajectories of both cerebral perfusion Satterthwaite et al. (2014), and structural covariance Váša et al. (2017). However, given that most regions showed approximately linear trajectories, we focused on results extracted using linear mixed effect models, and on linear rates of change ($\Delta FC_{14-26}$), in main analyses.

*Replication of analyses in an alternative parcellation*

As mentioned in the main text, all results and figures presented in the main text for the Human Connectome Project (Glasser et al., 2016) parcellation were repelicated with a different parcellation based on sub-parcellating the Desikan-Kiliany (DK) anatomical atlas (Desikan et al., 2006) into 308 parcels of approximately equal surface area ($\sim$5cm$^2$; Romero-Garcia et al., 2012). In Fig. S9, Fig. S10, Fig. S11 and Fig. S12 we show that the key results for all Figures in the main text hold in this alternative parcellation. Similarly, Table S4 recapitulates results from Table S3 with the DK sub-parcellation. Finally, Table S5 summarises the relationship between maturational index and a range of cortical maps in both parcellations.

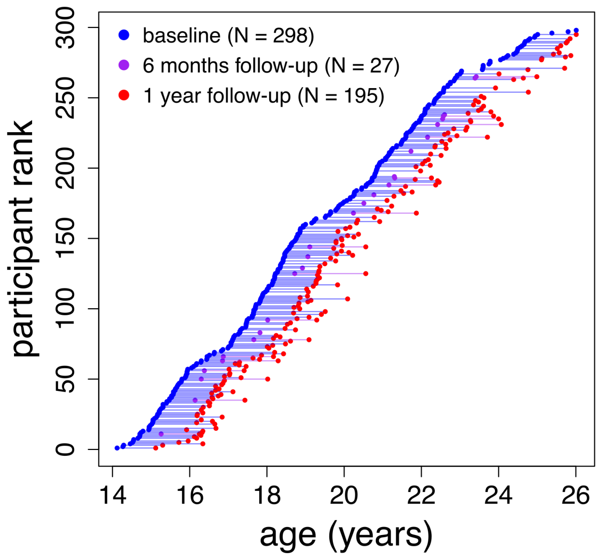


**Figure S1: Distribution of scans as a function of age.** Markers correspond to scans retained for inclusion in the study, connecting lines join consecutive scans for the same subject. Subjects are ordered by age at first "baseline" scan.

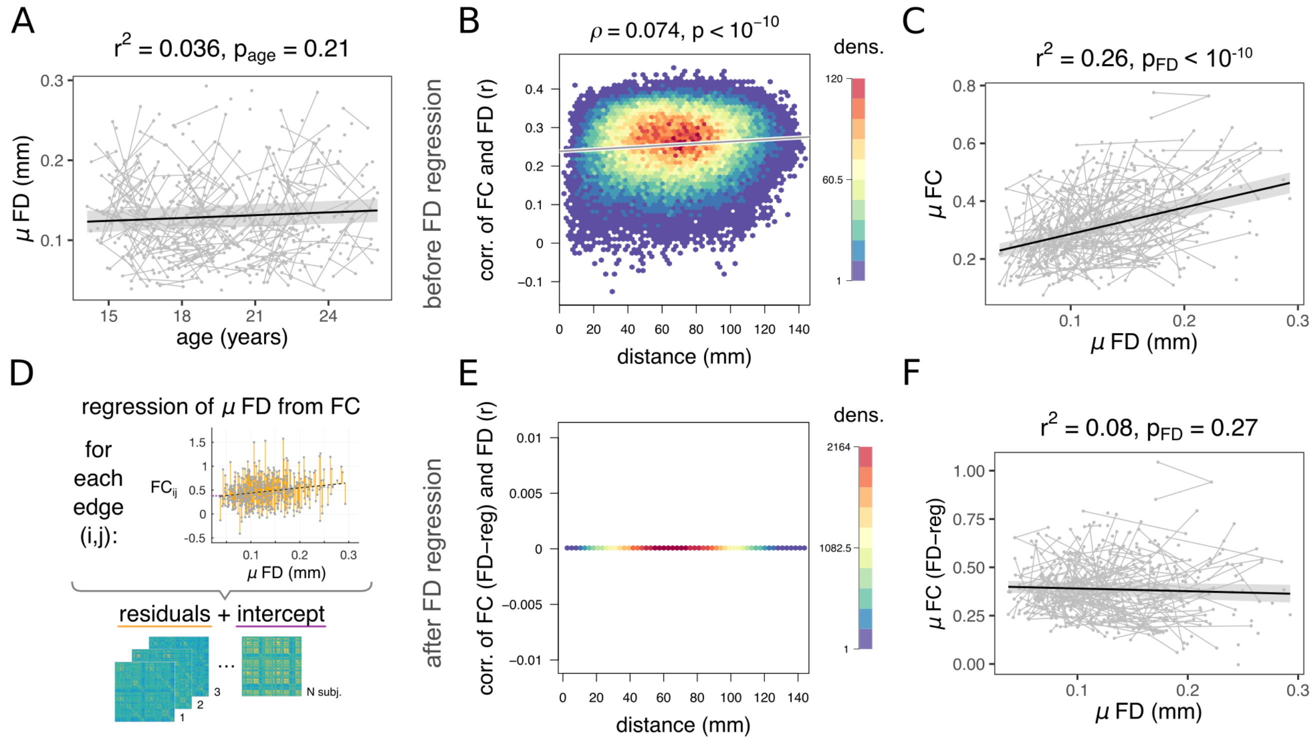


**Figure S2: Effects of head motion on functional connectivity.** A) Average participant motion (quantified as mean FD) does not change with age. B) The correlation between FC at each edge and the corresponding participant's motion (across participants) shows a weak relationship with the Euclidean distance spanned by edges. Moreover, the average edge-wise correlation between FC and motion (average of data along the y-axis) is non-zero. C) Mean participant motion is strongly related to mean FC across participants, such that the functional networks of subjects who move more are more strongly functionally connected. D) To remove the dependence of FC on motion, mean FD was regressed from each edge; the residuals constitute subject-specific FD-corrected FC, with intercepts retained to maintain the relative importance of edges across the group as well as the interpretability of FC values. Following edge-wise correction of FC for motion, the correlation between FC and motion vanished - both E) at the level of individual edges (by definition), and F) at the whole-brain level. Panels B and E depict 2D-histograms of the underlying dense data, with hexagonal bins color-coded by the number of edges located within.

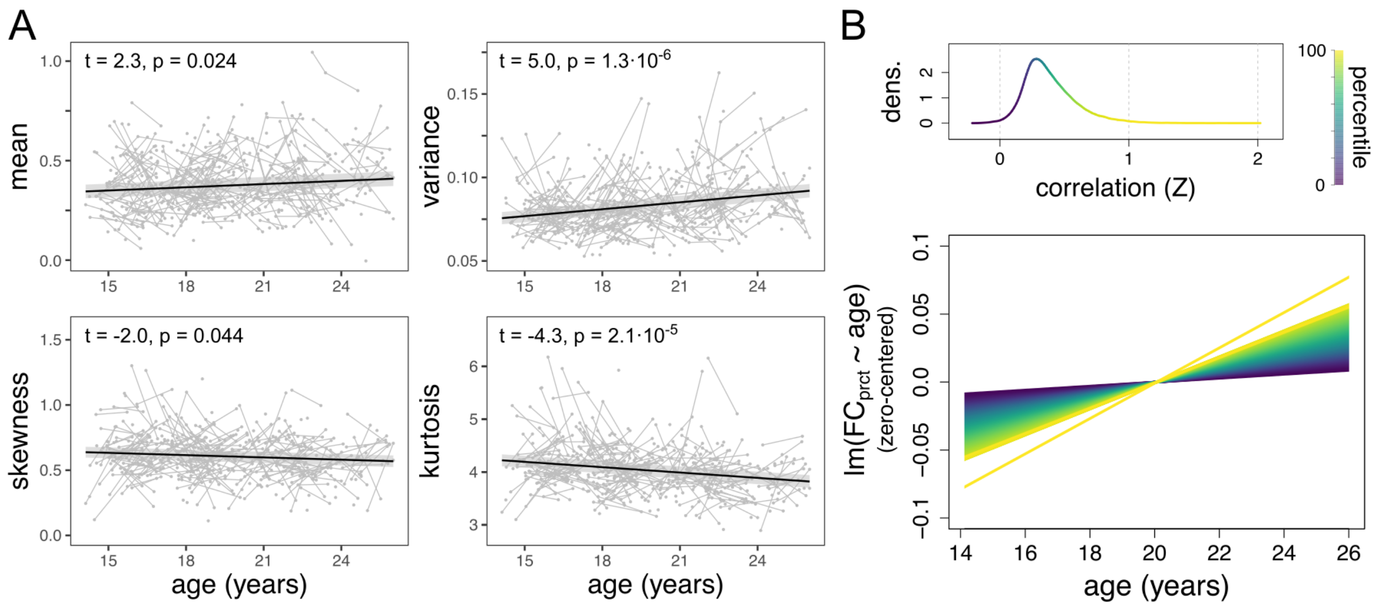


**Figure S3: Global trajectories of the correlation distribution.** A) Trajectories of the four moments of the correlation distribution as a function of age: (i) mean, (ii) variance, (iii) skewness and (iv) kurtosis. B) Example distribution, colour-coded by percentile. Higher percentiles of the correlation distribution strengthen faster, as indicated by increasing slopes of linear models fit to values of correlation at increasing percentiles. This is visualised using a plot of all the linear models as a function of age (centered along the y-axis for visual clarity).

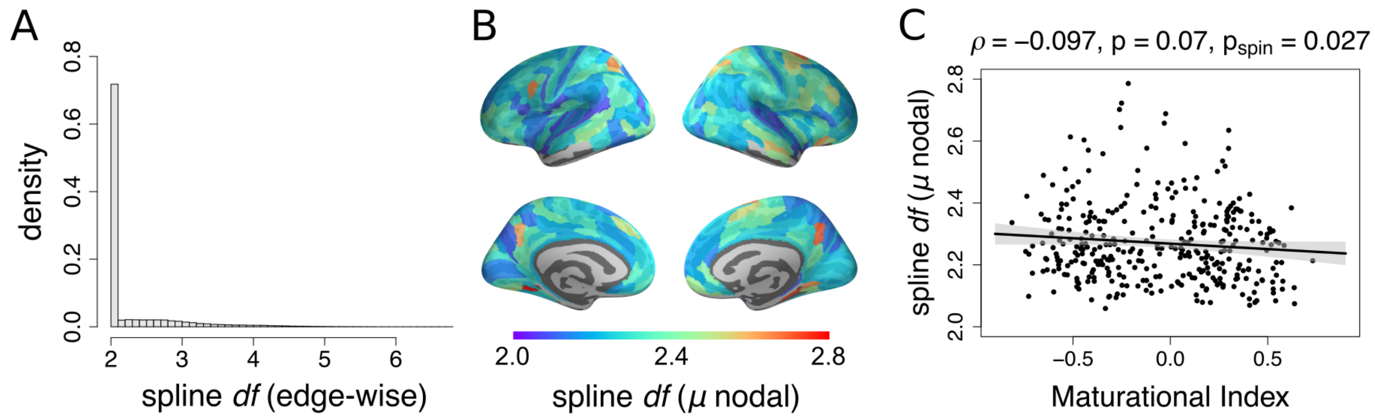


**Figure S4: Inspecting edge-wise trajectories for potential presence of non-linearities** A) Distribution of effective *df* of smoothing splines across edges. Most trajectories are linear: 71.7% trajectories had $df < 2.1$. B) Cortical distribution of average nodal *df* (averaged across all of a node's edges). C) Relationship of the maturational index (MI) with nodal *df*.

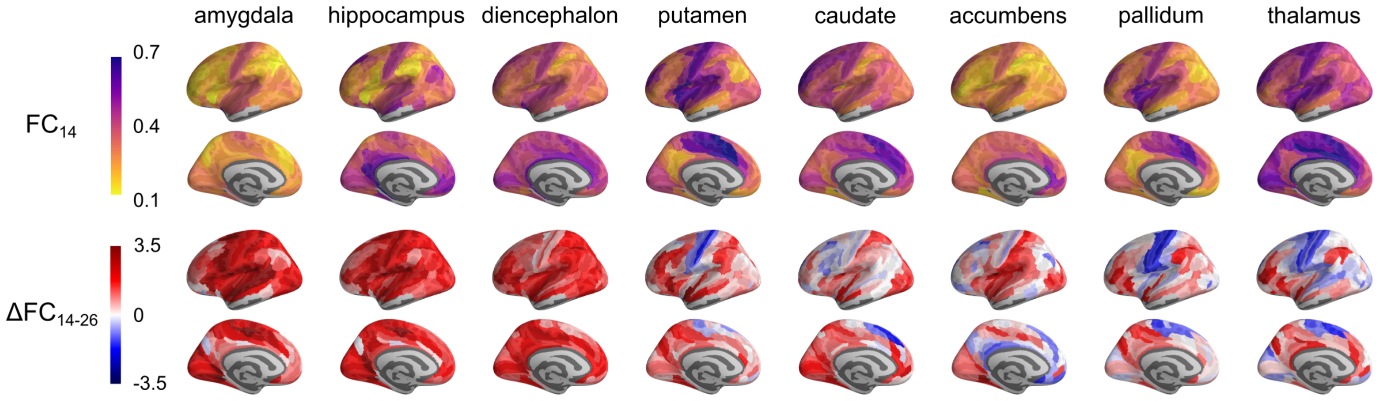


**Figure S5: Maturation of subcortico-cortical functional connectivity for all individual subcortical regions.** Parameters of linear models extracted from subcortico-cortical FC, averaged using the two edges between each bilateral pair of subcortical regions and each cortical region: $FC_{14}$ (top row) and $\Delta FC_{14-26}$ (bottom row). Maps are ordered from left to right in decreasing order of average rate of change with age. Due to space constraints as well as strong bilateral symmetry exhibited by the full maps, only the left hemisphere is visualised.

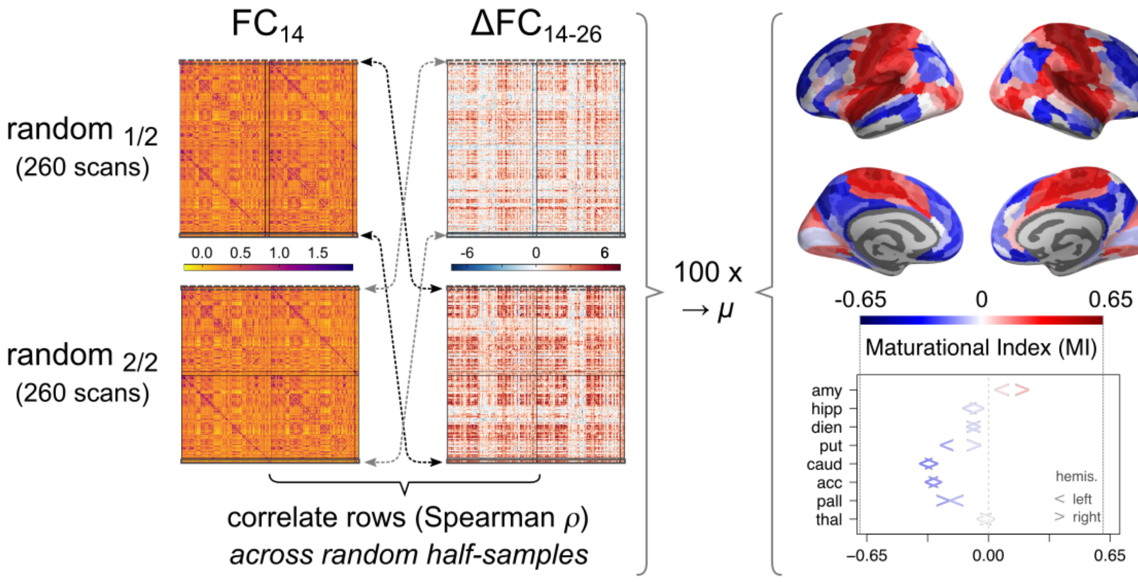


**Figure S6: Association cortical "disruptive" maturation is robust to random half-splitting of the data.** The set of 520 scans was randomly split in half, and edge-wise mixed-effect linear models were fitted separately to each half (of 260 scans), leading to two independent estimates of (i) FC at age 14 ($FC_{14}$), and (ii) the adolescent rate of change of FC ($\Delta FC_{14-26}$). The correlation between the two matrices was then calculated *across* the random half-samples, and averaged between the two split combinations. The results correspond to an average across 100 random half-splits.

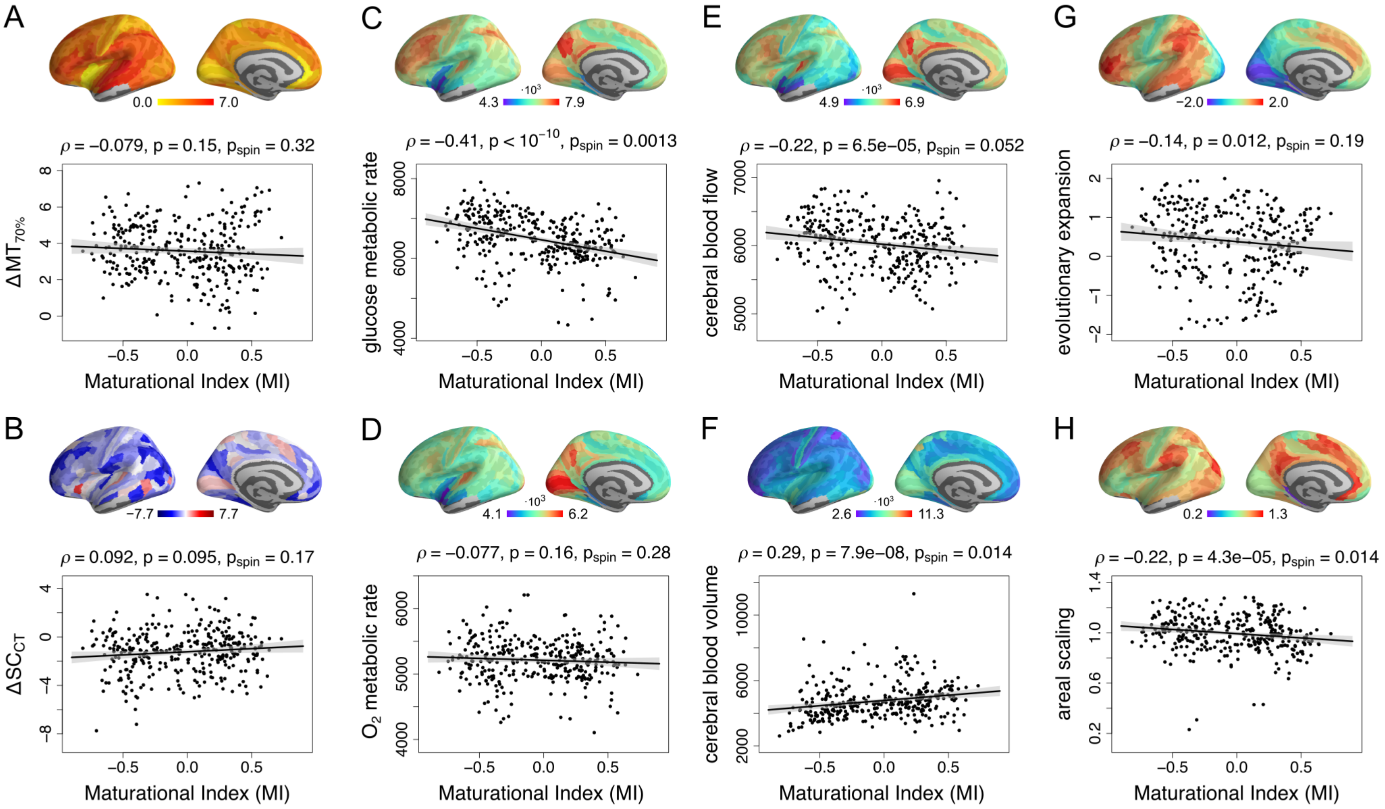


**Figure S7: Relationships between maturational index and additional cortical maps.** Spatial correlations between regional maturational index, and eight cortical maps: A) The rate of change of magnetization transfer (MT; a marker of myelination), extracted at 70% depth into cortex as in (Whitaker et al., 2016); B) The rate of change of nodal structural covariation of cortical thickness (CT) across subjects as in (Váša et al., 2017); C) metabolic rate of glucose; D) metabolic rate of oxygen; E) cerebral blood flow and F) cerebral blood volume, all (D-F) from Vaishnavi et al. (2010); G) rate of evolutionary expansion of the cortical surface, from Hill et al. (2010); and H) areal scaling of the cortical surface, from Reardon et al. (2018).

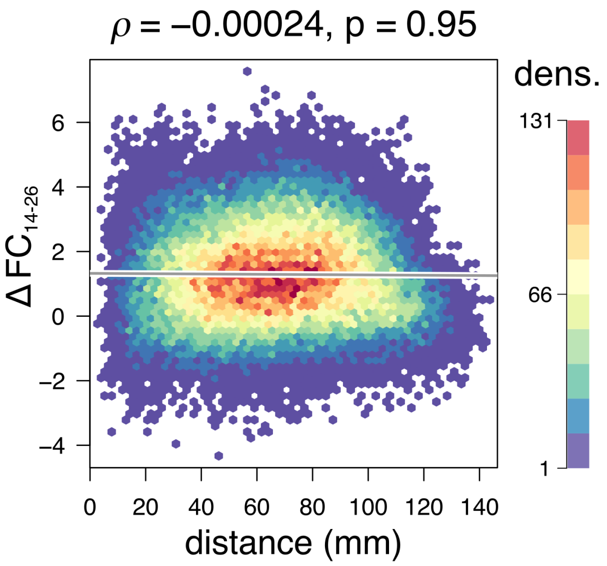


**Figure S8: Relationship of changes in FC to Euclidean distance.** Distances spanned by edges were calculated as the Euclidean distance between coordinates of centroid of the connected pair of regions. Changes in FC ($\Delta FC_{14-26}$) were extracted using mixed-effect linear models at the edge level, across participants. The plot is a 2D-histogram of the underlying dense data, with hexagonal bins color-coded by the number of edges located within.

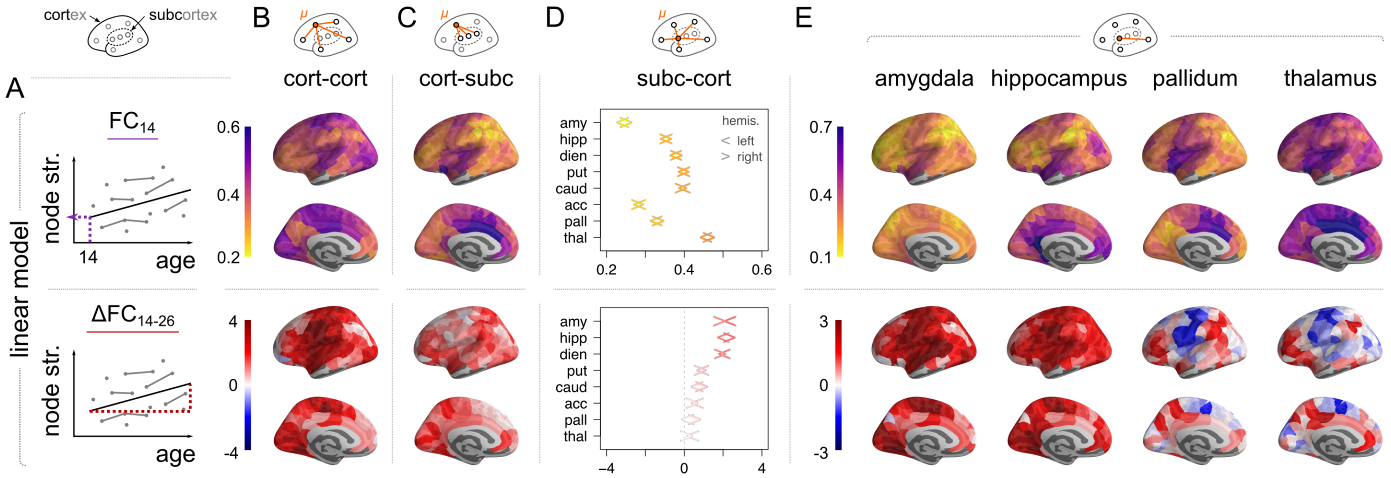


**Figure S9: Replication of main text Fig. 1 in the DK-sub parcellation: Regional strength of functional connectivity of cortical areas and subcortical nuclei at 14 years ($FC_{14}$) and regional change in strength of connectivity during adolescence ($\Delta FC_{14-26}$).** Regional strength for each of 282 cortical and 16 subcortical nodes was regressed on a linear function of age for all participants (N=298; mixed effects model). Connectivity at age 14 was strongest for primary motor and sensory cortical areas, thalamus and striatal regions. Adolescent change in cortico-cortical connectivity strength was also strongly positive for primary motor and sensory areas; but these areas had reduced strength of connectivity to subcortical nuclei, i.e., $\Delta FC_{14-26} < 0$. In contrast, connectivity between association cortex and subcortical nuclei, which was not strong at baseline, became more strongly positive during adolescence. In subcortico-cortical plots the left/right arrow corresponds to left/right hemisphere), and regions are ordered by average rate of change. Amygdala and hippocampus have weak cortical connectivity at baseline but the greatest rates of increase in cortical connectivity during adolescence. Thalamus has strong cortical connectivity at baseline but the greatest rates of decrease in primary cortical connectivity during adolescence.

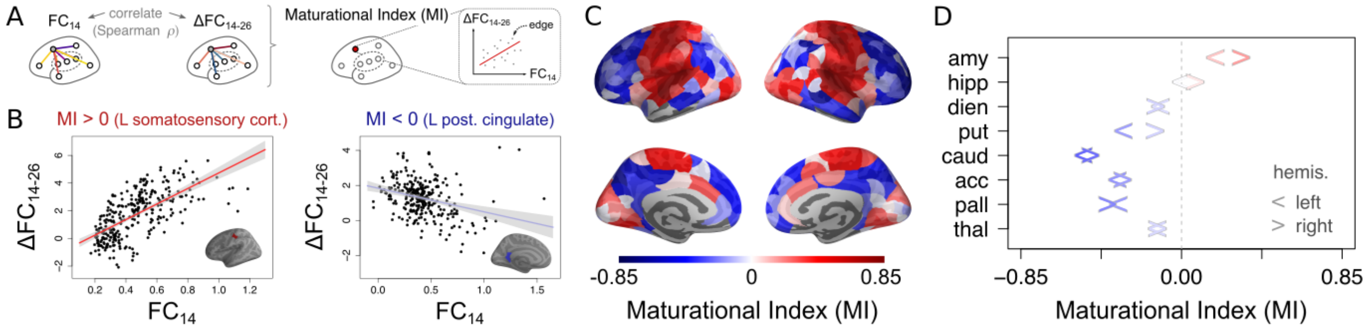


**Figure S10: Replication of main text Fig. 2 in the DK-sub parcellation: Maturational index.** A) The maturational index (MI) for each brain region is defined as the correlation of edge-wise baseline $FC_{14}$ versus rate of change $\Delta FC_{14-26}$. Panel B) illustrates this for two example regions, including a positive MI in left somatosensory cortex, and a negative MI in left posterior cingulate cortex. C) Visualisation of the Maturational Index for all cortical regions, and D) subcortical regions (the left/right arrow corresponds to left/right hemisphere).

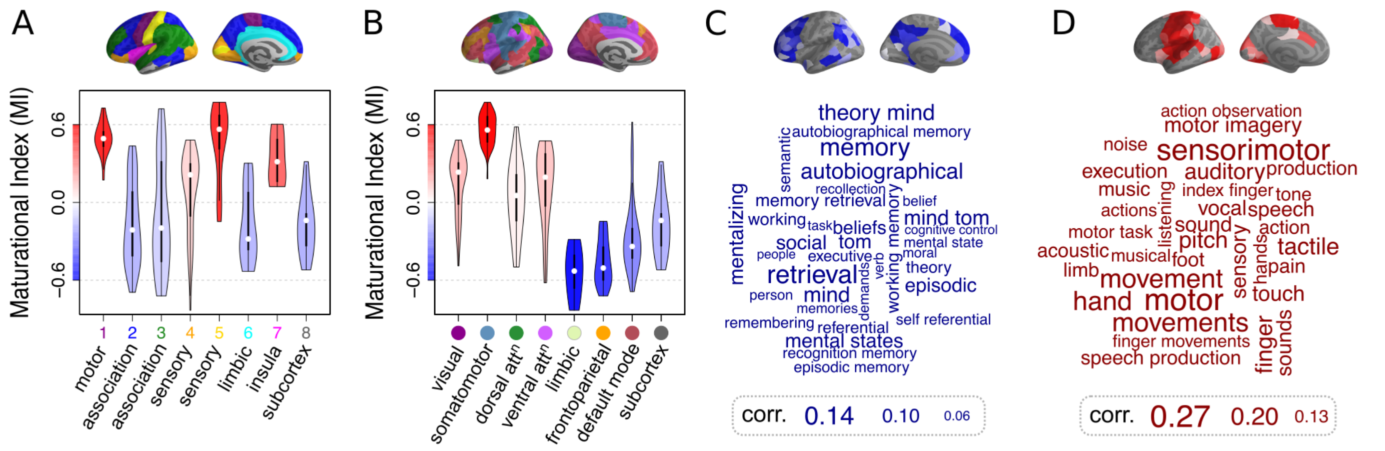


**Figure S11: Replication of main text Fig. 3 in DK-sub parcellation: Maturational index in anatomical and psychological context.** A) Distribution of maturational index for each cytoarchitectonic class of the von Economo atlas (Vértes et al., 2016), and B) for resting state networks derived from prior resting state FC analysis by Yeo (Yeo et al., 2011). In both cases, subcortical regions were considered as an additional eighth class/subnetwork. The violin plots are coloured by average MI within the corresponding class of regions. B) Word clouds of cognitive terms associated with cortical brain regions that have conservative (red) or disruptive (blue) modes of development (Neurosynth decoding (Yarkoni et al., 2011)). The size of cognitive terms corresponds to the correlation of corresponding meta-analytic maps generated by Neurosynth with each of the two modes (top).

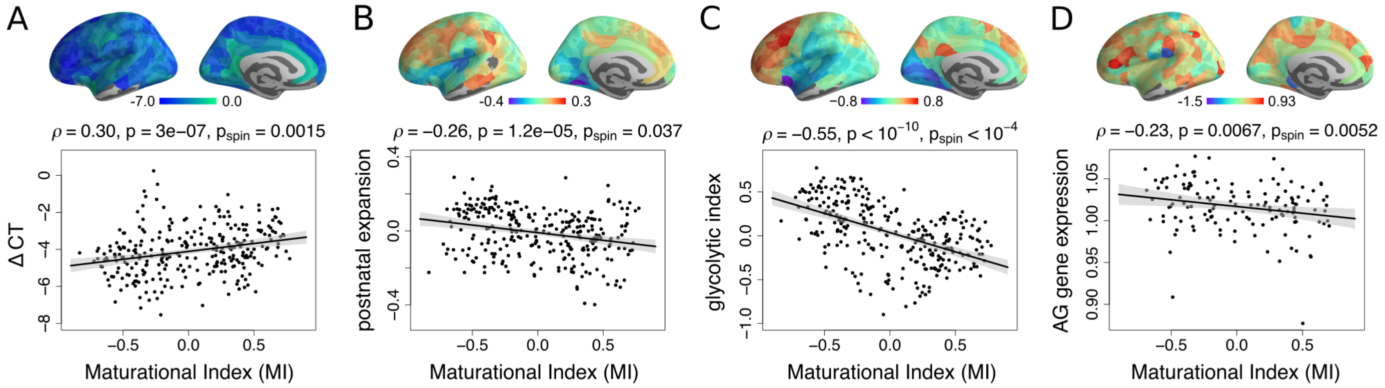


**Figure S12: Replication of main text Fig. 4 in the DK-sub parcellation: Disruptive and conservative modes of fMRI maturation in developmental and metabolic context.** A) Maturational index was positively correlated with $\Delta CT$ - regions which had disruptive development (MI<0) had faster rates of cortical thickness (CT) shrinkage during adolescence. B) MI was negatively correlated with a prior map of postnatal cortical surface area - disruptive maturation was greater in regions that showed greatest expansion after birth. C) MI was negatively correlated with a prior map of the glycolytic index, a measure of aerobic glycolysis (AG); and D) MI was negatively correlated with a prior map of brain regional expression of AG-related genes. (Hill et al., 2010).

| **Paper** | **Age (y)** | **# Sub.** | **Motion** $\star$ | **GSR** | **Nodes** | **Edges** | **Thresh.** | **Results** |
|---|---|---|---|---|---|---|---|---|
| Fair<br>PNAS<br>2007 | 7-31 | 139 | motion match | yes | 39 | Pears. r | r $\geq$ 0.1 | • sub-networks segregate with age<br>• long edges increase in strength, short edges decrease |
| Fair<br>PLoS CB<br>2009 | 7-31 | 210 | none | yes | 34 | Pears. r | r $\geq$ 0.1 | • shift from 'local' to 'distributed' communities<br>• community assignment changes; modularity stable<br>• path length, clustering, small-worldness stable |
| Supekar<br>PLoS CB<br>2009 | 7-22 | 45 | none | no | 90 | wav. corr. | binary 1-99% | • path length ($\lambda$), efficiency, clustering ($\gamma$), SWI $\sigma$ stable<br>• paralimbic weights incr., subcortical weights decr.<br>• long edges increase, short edges decrease |
| Stevens<br>HBM<br>2009 | 12-30 | 100 | none | no | voxels | Grang. caus. | N/A | • linear (or log) decrease in causal interactions, esp. betw. lateral prefrontal-parietal circuits and DMN |
| Dosenb.<br>Science<br>2010 | 7-30 | 238 +195 | none | yes | 160 | Pears. r | none | • strengthening edges longer than weakening edges<br>• within-module edges strengthen, betw.-module weaken<br>• MVPA explains 55% variance of changes with age |
| Anderson<br>Brain Con.<br>2011 | 7-35 | 1278 | none | no | 7266 | Pears. r | none | • connectivity decrease betw. attention control and DMN<br>• DMN "boundaries sharpen" (conn. within incr.)<br>• slight decr.in connectivity within frontoparietal ROIs |
| Satterthw.<br>NeuroIm.<br>2013 | 8-22 | 780 | motion match | • yes<br>• no<br>(both) | 264 | Pears. r | none | • motion inflates effects of age & distance-dependence<br>• motion dampens incr. in intra-modular connectivity<br>+ large N needed to detect changes in conn. w/ age |
| Hwang<br>Cer. Cor.<br>2013 | 10-20 | 99 | time despike | no | • voxels<br>• 160 | Pears. r | mult. wei. + bin. | • hub architecture stable<br>• early incr. betw. frontal hubs & cort. + subc. non-hubs<br>• late incr. betw. cerebellar hubs & cortical non-hubs |
| Wu<br>PLoS One<br>2013 | 5.7-18.4 | 60 | none | yes | 90 | Pears. r | binary 20-35% | • path length (PL, $\lambda$), efficiency, clust. (cc) stable<br>• clustering ($\gamma$), SWI $\sigma$, modularity Q increase<br>• nodal changes in degree, efficiency and betw. cent. |
| Grayson<br>PLoS One<br>2014 | 7-35 | 21 | scrub. | yes | 219 | Pears. r | binary 1-10% | • rich club strengthening (rich $\rightarrow$ richer & poor $\rightarrow$ rich)<br>• strengthening edges: somatosensory, insula (NBS)<br>+ structural (DWI) rich club organisation stable |
| Satterthw.<br>PNAS<br>2014 $*$ | 8-22 | 922 | covaried | N/A | voxels | Pears. r | N/A | • perfusion decr. nonlinear (min. $\sim$17y), esp. assoc. cortex<br>+ trajectory diff. betw. females (decr. U) and males ($\sim$lin.) (sex differences appear only mid-puberty) |

| Paper | Age (y) | # Sub. | Motion ⋆ | GSR | Nodes | Edges | Thresh. | Results |
|---|---|---|---|---|---|---|---|---|
| Gu<br>PNAS<br>2015 | 8-22 | 780 | motion regress. from mat. | no, but mat. norm. by $\mu$ | 264 | Pears. r | none | • variance of module size & similarity to adult modules incr.<br>• DMN: within + between-module strength increases<br>• hi.-order, subc./cerebell.: w/in + betw.-mod. strength decr. |
| Marek<br>PLoS CB<br>2015 | 10-26 | 192 | wavelet despike + scrub. | no | 264 | Pears. r | binary 1-25% | • module organization stable<br>• cing.-operc./salience part. coeff. incr. ($\mapsto$ inhibitory control)<br>• no distance-dependent change of strength |
| Kaufmann<br>Nat. Neurosci.<br>2017 | 8-22 | 797 | AROMA + FIX | no | 227 = 268 -41 drop. | Pears. r | weighted | • resting-state, working memory task, emotion recognition task<br>• "fingerprinting" used to identify individual across scans<br>• identification accuracy ("distinctiveness") of indiv. FC incr. |
| Kundu<br>J. Neurosci.<br>2018 | 8.3-46.2 | 51 | ME-ICA | no | • voxels<br>• D-K | Pears. r | weighted r=0.3–0.9 | • n° BOLD components decreases exponentially<br>• driven by dlPFC, parietal cortex, cerebellum<br>• BOLD dimensionality correlates to part. coeff. |

**Table S1:** Summary of adolescent development of whole-brain resting-state functional networks. Only studies whose age-range presents overlap with the NSPN study of adolescent development (ages 14-24 years) were selected; studies on "lifespan" development whose age-range extends up to old age were not included. ⋆ Methods for motion correction listed in the "Motion" column do not include linear regression of motion parameters and their derivatives (which was applied by most studies). ∗ The study by Satterthwaite et al. (2014) (listed as Satterthw. PNAS 2014) is an arterial spin labeling (ASL) study.

| Paper | Age (y) | # Sub. | Motion ⋆ | GSR | Edges | hip | amy | die | pal | put | acc | cau | thal |
|---|---|---|---|---|---|---|---|---|---|---|---|---|---|
| Qin PNAS 2012 | 7-22 | 87 | none | yes | mult. regress$^{n}$ | | ↑ para+limb. ass$^{n}$ vmPFC | | | | | | |
| Gabard-D. NeuroIm. 2014 | 4-23 | 58 | scrub. + FD reg. | yes | mult. regress$^{n}$ | | ↑ mPFC ↓ ins. / STS parah. / PCC | | | | | | |
| Greene J. Neurosci. 2014 | 7-31 | 180 | scrub. | no | partial corr. | | | | ↓ somatomotor face system (BG: pal. + put.) | | | | |
| Alarcon NeuroIm. 2015 | 10-16 | 122 | scrub. | yes | Pears. r | | ↓ par.-occip. ↓ mFC | | | | | | |
| Fareri NeuroIm. 2015 | 4.5-23 | 66 | scrub. + FD reg. | yes | mult. regress$^{n}$ | | | | | | ventral striatum (put. + acc. + cau.) ↓ mPFC | | |
| Sato NeuroIm. 2015 | 7-15 | 447 | scrub. | yes | Pears. r | | | | | | | ↓ e-vec. cent. | ↓ e-vec. cent. |
| van Duij. NeuroIm. 2015 | 8-25 | 299 | FD reg. | yes | dual regress$^{n}$ | ↑ nucl. acc. | | | | | ↑ dACC hipp. | ↑ dlPFC | ↑ dlPFC |

**Table S2:** Summary of adolescent development of subcortical resting-state functional networks. Only studies whose age-range presents substantial overlap with the NSPN study of adolescent development (ages 14-24 years) were selected; studies on "lifespan" development whose age-range extends up to old age were not included. ⋆ Methods for motion correction listed in the "Motion" column do not include linear regression of motion parameters and their derivatives (which was applied by most studies).

**Table S3:** Maturation of subcortico-cortical FC. Frequencies of increases (↑) and decreases (↓) in subcortico-cortical FC as a function of age. These frequencies are listed as overall (all), and as numbers satisfying $P < 0.05$ - both without ($P_{\text{raw}}$) and with ($P_{\text{FDR}}$) correction of P-values for multiple comparisons. Subcortical regions are listed in decreasing order of average rate of change with age.

| | | amygdala | hippocampus | diencephalon | putamen | caudate | accumbens | pallidum | thalamus |
|---|---|---|---|---|---|---|---|---|---|
| | all | 327 | 333 | 333 | 278 | 277 | 231 | 230 | 206 |
| ↑ | $P_{\text{raw}}$ | 161 | 119 | 99 | 26 | 13 | 24 | 8 | 18 |
| | $P_{\text{FDR}}$ | 69 | 5 | 0 | 0 | 0 | 0 | 0 | 0 |
| | all | 19 | 13 | 13 | 68 | 69 | 115 | 116 | 140 |
| ↓ | $P_{\text{raw}}$ | 0 | 0 | 0 | 0 | 2 | 2 | 2 | 0 |
| | $P_{\text{FDR}}$ | 0 | 0 | 0 | 0 | 0 | 0 | 0 | 0 |

**Table S4:** Maturation of subcortico-cortical FC, in the DK-sub parcellation. Frequencies of increases (↑) and decreases (↓) in subcortico-cortical FC as a function of age. These frequencies are listed as overall (all), and as numbers satisfying $P < 0.05$ - both without ($P_{\text{raw}}$) and with ($P_{\text{FDR}}$) correction of P-values for multiple comparisons. Subcortical regions are listed in decreasing order of average rate of change with age.

| | | amygdala | hippocampus | diencephalon | putamen | caudate | accumbens | pallidum | thalamus |
|---|---|---|---|---|---|---|---|---|---|
| | all | 278 | 287 | 289 | 230 | 238 | 203 | 193 | 163 |
| ↑ | $P_{\text{raw}}$ | 132 | 106 | 71 | 14 | 10 | 11 | 2 | 13 |
| | $P_{\text{FDR}}$ | 43 | 0 | 0 | 0 | 0 | 0 | 0 | 0 |
| | all | 20 | 11 | 9 | 68 | 60 | 95 | 105 | 135 |
| ↓ | $P_{\text{raw}}$ | 0 | 0 | 0 | 0 | 0 | 2 | 1 | 0 |
| | $P_{\text{FDR}}$ | 0 | 0 | 0 | 0 | 0 | 0 | 0 | 0 |

**Table S5:** Relationships of maturational index with a range of cortical maps. Map names on the left correspond to maps visualised in main text Fig. 4, and in supplementary Fig. S7 and S12. Columns contain the Spearman $\rho$, corresponding $P$-value as well as the $P_{\text{spin}}$ $P$-value generated using the spatial permutation test - for both the HCP and DK-sub parcellations.

| parcellation → | HCP | | | DK-sub | | | |
|---|---|---|---|---|---|---|---|
| map name ↓ | Spear. $\rho$ | $P$ | $P_{\text{spin}}$ | Spear. $\rho$ | $P$ | $P_{\text{spin}}$ | reference ↓ |
| Δ CT | 0.16 | 0.0033 | 0.026 | 0.30 | $3.0{\cdot}10^{-7}$ | 0.0015 | Whitaker et al., 2016 |
| Δ $MT_{70\%}$ | -0.079 | 0.15 | 0.32 | -0.14 | 0.018 | 0.18 | Whitaker et al., 2016 |
| Δ $SC_{CT}$ | 0.092 | 0.095 | 0.17 | 0.031 | 0.60 | 0.38 | Váša et al., 2017 |
| glucose metabolic rate | -0.41 | $< 10^{-10}$ | 0.0013 | -0.33 | $1.6{\cdot}10^{-8}$ | 0.008 | Vaishnavi et al., 2010 |
| glycolytic index | -0.56 | $< 10^{-10}$ | $< 10^{-4}$ | -0.55 | $< 10^{-10}$ | $< 10^{-4}$ | Vaishnavi et al., 2010 |
| $O_2$ metabolic rate | -0.077 | 0.16 | 0.28 | 0.057 | 0.35 | 0.36 | Vaishnavi et al., 2010 |
| cerebral blood volume | 0.29 | $7.9{\cdot}10^{-8}$ | 0.014 | 0.24 | $5.9{\cdot}10^{-5}$ | 0.052 | Vaishnavi et al., 2010 |
| cerebral blood flow | -0.22 | $6.5{\cdot}10^{-5}$ | 0.052 | -0.16 | 0.0078 | 0.11 | Vaishnavi et al., 2010 |
| postnatal expansion | -0.28 | $1.9{\cdot}10^{-7}$ | 0.023 | -0.26 | $1.2{\cdot}10^{-5}$ | 0.037 | Hill et al., 2010 |
| evolutionary expansion | -0.14 | 0.012 | 0.19 | -0.26 | $9.9{\cdot}10^{-6}$ | 0.025 | Hill et al., 2010 |
| areal scaling | -0.22 | $4.3{\cdot}10^{-5}$ | 0.014 | -0.18 | 0.0027 | 0.037 | Reardon et al., 2018 |
| AG gene expression | -0.34 | $6.1{\cdot}10^{-6}$ | $< 10^{-4}$ | -0.23 | 0.0067 | 0.0052 | Hawrylycz et al., 2012 |
